# SARS-ANI: A Global Open Access Dataset of Reported SARS-CoV-2 Events in Animals

**DOI:** 10.1101/2022.04.11.487836

**Authors:** Afra Nerpel, Liuhuaying Yang, Johannes Sorger, Annemarie Käsbohrer, Chris Walzer, Amélie Desvars-Larrive

**Affiliations:** Unit of Veterinary Public Health and Epidemiology, University of Veterinary Medicine Vienna, Veterinaerplatz 1, 1210 Vienna, Austria; Complexity Science Hub Vienna, Josefstaedter Strasse 39, 1080 Vienna, Austria; Wildlife Conservation Society, 2300 Southern Blvd, Bronx, NY 10460, United States; Research Institute of Wildlife Ecology, University of Veterinary Medicine Vienna, Savoyenstrasse 1, 1160 Vienna, Austria; VetFarm, University of Veterinary Medicine Vienna, Kremesberg 13, 2563 Pottenstein, Austria

**Author notes:** Corresponding author: Amélie Desvars-Larrive.

## Abstract

The zoonotic origin of SARS-CoV-2, the etiological agent of COVID-19, is not yet fully resolved. Although natural infections in animals are reported in a wide range of species, large knowledge and data gaps remain regarding SARS-CoV-2 animal hosts. We used two major health databases to extract unstructured data and generated a comprehensive global dataset of thoroughly documented SARS-CoV-2 events in animals. The dataset integrates relevant epidemiological and clinical data on each event and is readily usable for analytical purposes. We also share the code for technical and visual validation of the data and created a user-friendly dashboard for data exploration. Data on SARS-CoV-2 occurrence in animals is critical to adapt monitoring strategy, prevent the formation of animal reservoirs, and tailor future human and animal vaccination programs. The FAIRness and analytical flexibility of the data will support research efforts on SARS-CoV-2 at the human-animal-environment interface. We intend to update this dataset weekly for at least one year and, through collaborative processes, to develop the dataset further and expand its use.

## Background & Summary

Although the coronavirus disease 2019 (COVID-19) pandemic, caused by the severe acute respiratory syndrome coronavirus 2 (SARS-CoV-2), is driven by human-to-human transmission, emergence in humans most likely involved at least two independent zoonotic spillover (i.e. animal-to-human transmission) events from wild animals kept at the Huanan Seafood Market in Wuhan, China^1,2^. The first officially confirmed case in an animal was reported in February 2020, when a dog in Hong Kong tested positive for the virus shortly after its owner was diagnosed with COVID-19^3^. SARS-CoV-2 is recognized as a generalist coronavirus^4^, showing a great capacity for infecting multiple animal species^5^, including pets, e.g. dogs^3,6,7^, cats^6,8,9^, and Syrian hamsters^10^; zoo animals^11^, e.g. gorillas (https://www.science.org/content/article/captive-gorillas-test-positive-coronavirus), lions^12–14^, tigers^12,14^, mountain lion^13^, and Asian small-clawed otter (https://www.aphis.usda.gov/aphis/newsroom/stakeholder-info/sa_by_date/sa-2021/sa-04/covid-georgia-otters); farmed animals, e.g. mink^15–17^; and free-ranging wildlife^11^, e.g. white-tailed deer^18–20^ and leopard^21^. SARS-CoV-2-infected animals may experience subclinical to severe signs of infection^3,14,17^ and implemented interventions range from individual treatment to preventive culling. Animal infections mostly result from human-to-animal transmission (“spillback”) and can, in certain cases, lead to onward epizootic circulation of the virus, among conspecifics, e.g. in hamsters^22^, mink^16^, tigers^23^, and white-tailed deer^18–20^, or between species, e.g. from mink to cats^24^. Additional spillback events from humans to animals may have occurred in the past two years although as yet undetected^25^. Recently, animal-to-human transmissions were observed from farmed mink^16^, pet hamster^10^, and possibly free-ranging white-tailed deer^26^. These secondary spillovers resulted in the mass culling of mink^27^ and occasionally led to subsequent human-to-human transmission^10^. Not only does SARS-CoV-2 represent a risk for public health, but it is also a threat for animal health and welfare as well as conservation^28,29^.

Reports on SARS-CoV-2 in animals are primarily available from the Program for Monitoring Emerging Diseases (ProMED-mail) (https://promedmail.org/) and the World Organisation for Animal Health (OIE) World Animal Health Information System (WAHIS) (https://wahis.oie.int/). However, this data is unstructured (narrative text) and/or available in multiple excel sheets or PDF files, therefore not usable without preliminary, time-consuming curation and formatting steps. The Animal and Plant Health Inspection Service (APHIS) of the U.S. Department of Agriculture (USDA) publishes a dashboard of confirmed cases of SARS-CoV-2 in animals in the United States (https://www.aphis.usda.gov/aphis/dashboards/tableau/sars-dashboard) while the Canadian Animal Health Surveillance System (CAHSS) has a dashboard for Canada (https://cahss.ca/cahss-tools/sars-cov-2-dashboard). Data can be downloaded from the APHIS-USDA dashboard, but only as an image, PDF, or PowerPoint (not machine-readable) whereas underlying data are not accessible from the CAHSS dashboard. Both the Danish and Dutch governments maintain a website dedicated to SARS-CoV-2 in mink farms in the respective country (https://www.foedevarestyrelsen.dk/Dyr/Dyr-og-Covid-19/Mink-og-COVID-19/Sider/Kort-over-kommuner-med-smittede-minkfarme.aspx and https://www.rijksoverheid.nl/actueel/nieuws?trefwoord=SARS-CoV-2&startdatum=11-07-2020&einddatum=18-09-2020&onderdeel=Alle+ministeries, respectively), without any possibility of accessing the raw data.

Although the OIE, Food and Agriculture Organization (FAO), and World Health Organization (WHO) recently published a statement on “*the prioritization of monitoring SARS-CoV-2 infection in wildlife and preventing the formation of animal reservoirs*” (https://www.who.int/news/item/07-03-2022-joint-statement-on-the-prioritization-of-monitoring-sars-cov-2-infection-in-wildlife-and-preventing-the-formation-of-animal-reservoirs), there is currently no comprehensive global dataset on SARS-CoV-2 events in animals that can be easily imported, processed, and analysed.

In the context of the current COVID-19 pandemic, availability of FAIR (Findable, Accessible, Interoperable, and Reusable) data^30^ is critical to understand the current and developing epidemiology of SARS-CoV-2 at human-animal interfaces and mitigate the impacts of this and future pandemics. In this paper, following the Open Science Principles^31,32^, we document and share the methods used to produce *SARS-ANI*, a comprehensive curated global dataset of thoroughly documented SARS-CoV-2 events in animals. We provide a detailed description of the dataset and supplement it with user-friendly documentation and materials (code and archived reports) to enhance data comprehension and use. We also present examples of usages to provide insights into the epidemiological and clinical patterns of SARS-CoV-2 animal infections globally, at the country and species level. The generated dataset is publicly available and readily usable for analytical purposes. The SARS-ANI Dataset and its dashboard greatly facilitate access to and re-use of data on SARS-CoV-2 events in animals. The dashboard also allows non-experts to access and view SARS-CoV-2 animal events. The continuous analysis of SARS-CoV-2 occurrence data in animals is especially critical to adapt monitoring, surveillance and vaccination programs for animals and humans in a timely manner and evaluate the developing threat SARS-CoV-2 represents for public and animal health as well as biodiversity and conservation.

## Methods

Data for this dataset was collected from two major animal health databases: the Program for Monitoring Emerging Diseases (ProMED-mail) (https://promedmail.org/) and the World Organisation for Animal Health-World Animal Health Information System (OIE-WAHIS) (https://wahis.oie.int/).

### Step 1: Incorporating ProMED-mail Reports

ProMED-mail (https://promedmail.org/) is the largest publicly available system reporting global infectious disease outbreaks (outbreak denotes the occurrence of one or more cases in an epidemiological unit). It provides reports (called “posts”) on outbreaks and disease emergence. The information flow leading to publication of ProMED-mail reports is as follows: a disease event to be dispatched is selected from daily notifications of outbreaks received via emails, searching through the Internet and traditional media, and scanning of official and unofficial websites. All incoming information is reviewed and filtered by an editor or associate editor who, subsequently sends them to a multidisciplinary team of subject matter expert moderators who assess the accountability and accuracy of the information, interpret it, provide commentary, and give references to previous ProMED-mail reports and the scientific literature^33^. One ProMED-mail report, identified via a unique report identifier, may depict one single or several health events.

The incorporation of the ProMED-mail reports of interest followed two steps:

i. Selection of ProMED-mail reports We identified reports describing SARS-CoV-2 events in animals, i.e. presenting at least one individual case of SARS-CoV-2 infection or exposure in an animal, via the “Search Posts” function provided on the ProMED-mail website. We used the keywords “animal” and “COVID-19”, which are consistently used in the “Subject” of the ProMED-mail posts to report information related to SARS-CoV-2 in animals, to retrieve the reports pertaining to natural and experimental infections or vaccine assays in animals as well as general discussions on SARS-CoV-2 in animals (note: although COVID-19 refers to the disease caused by SARS-CoV-2 in humans and should not be used for animals, ProMED-mail conveniently uses this keyword for both humans and animals). Reports describing naturally occurring infection (i.e. the presence of the virus is demonstrated through laboratory method(s)) or exposure (i.e. the presence of antibodies against SARS-CoV-2 is evidenced through laboratory method(s)) of a single individual or group of individuals were filtered manually and considered for data extraction. As of date of submission (11 April 2022), the ProMED-mail database included 231 reports on SARS-CoV-2 in animals.
ii. Link to previous reports When a health event is continuing, ProMED-mail publishes follow-up reports, which provide references to previous ProMED-mail reports (at the end of the report or in the section “See Also” at the end of the post). We used this information to identify potential relationship of each reported event to a previous one (e.g. clinical follow-up, further spread of the virus, and treatment outcome) and entered this data into the final dataset.

### Step 2: Incorporating OIE-WAHIS Reports

OIE-WAHIS (https://wahis.oie.int/) is a Web-based computer system that processes data on animal diseases in real-time. OIE-WAHIS data reflects the information gathered by the Veterinary Services from OIE Members and non-Members Countries and Territories on OIE-listed diseases in domestic animals and wildlife, as well as on emerging and zoonotic diseases. In accordance with the OIE Terrestrial Animal Health Code^34,35^, the detection of infection with SARS-CoV-2 in animals meets the criteria for reporting to the World Animal Health Organisation (OIE) as an emerging infection. Only authorised users, i.e. the Delegates of OIE Member Countries and their authorised representatives, can enter data into the OIE-WAHIS (https://wahis.oie.int/) platform to notify the OIE of relevant animal disease information.

One OIE-WAHIS report, denominated via a unique report identifier, may contain one single or several outbreaks, each identified via a unique outbreak identifier. All information are publicly accessible on the OIE-WAHIS (https://wahis.oie.int/) interface.

The incorporation of the OIE-WAHIS reports of interest was performed in two steps:

i. Selection of OIE-WAHIS reports We used the OIE-WAHIS Dashboard of animal health events (https://wahis.oie.int/#/events) to extract cases of SARS-CoV-2 infection in animals notified by OIE Member and non-Members States. OIE-WAHIS publishes immediate notifications (INs) and follow-up reports (FURs), identifiable through the prefix “IN” and “FUR” in their respective names. Immediate notifications dispense information on newly notified events while FURs generally provide updates on previously notified, ongoing events (e.g. number of newly infected animals and new deaths, newly implemented control measures). We applied filters to the field “DISEASE” (“SARS-CoV-2 in animals (inf. with)”) and “REPORT DATE” to select reports related to SARS-CoV-2 events from 1^rst^ December 2019 until today. The reports can be consulted online or downloaded as an individual PDF or Excel file, each file corresponding to one country report (i.e. several outbreaks can be included in one report). As of date of submission (11 April 2022), the OIE-WAHIS Dashboard included 294 reports on SARS-CoV-2.
ii. Identification of gaps and dataset completion ProMED-mail screens a large range of information sources including OIE-WAHIS reports. The ProMED-mail posts mention the event ID of the OIE-WAHIS report(s) used as information source, which makes it possible to consult the original source on the OIE-WAHIS Dashboard (https://wahis.oie.int/#/events). OIE-WAHIS in many cases provides more succinct, technical reports. Therefore, we chose to first identify SARS-CoV-2 events in animals in the ProMED-mail database. In a second step, we used the OIE-WAHIS Dashboard (https://wahis.oie.int/#/events) to identify gaps, i.e. complete the previously entered SARS-CoV-2 events (hereinafter referred to as *sibling events*) and find additional events not reported in ProMED-mail (Fig. 1). For each country (using the filter “COUNTRY/TERRITORY” on the OIE-WAHIS Dashboard), we identified sibling events by comparing the OIE-WAHIS reports against all the previously entered ProMED-mail reports of the country, using the information on species, subnational administration, and date of laboratory confirmation (a buffer of ± 7 days was considered due to possible discrepancies related to confirmation by different laboratories) or date of publication when date of laboratory confirmation was missing (in this case a buffer of 30 days was considered because date of publication is strongly database-dependent). We did not use information about the city here because reports may inconsistently refer to city/village of outbreak occurrence due to data privacy reason. This strategy, although time consuming, was consistently applied throughout the data extraction process, ensuring a comprehensive collection of information for each outbreak, accuracy of the data, and reproducibility of the method.

**Fig. 1.**
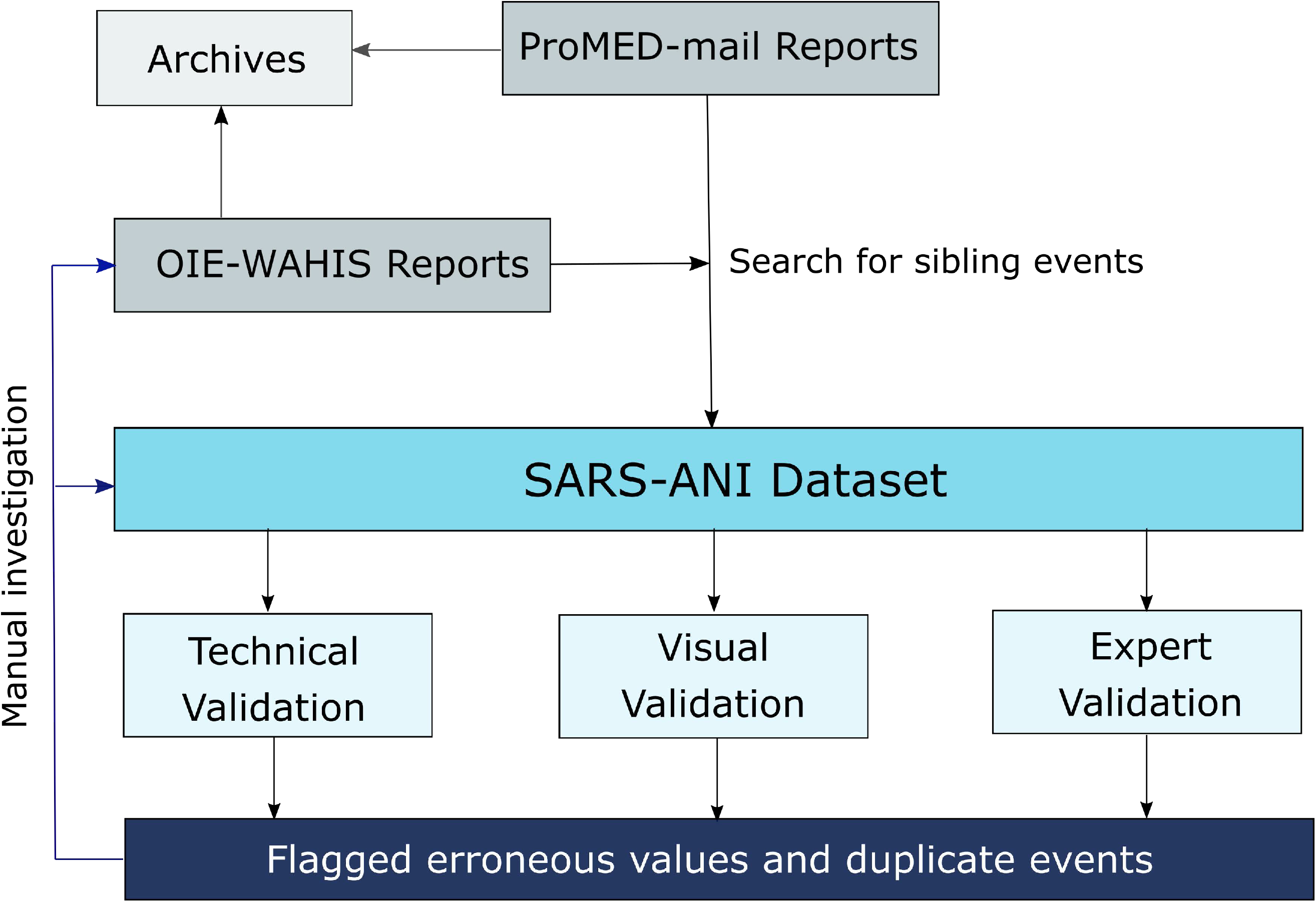
Schematic overview of the methodology: report integration and validation steps.

### Data Extraction

ProMED-mail provides detailed, text-based (narrative) reports of health events. This data is unstructured whereas OIE-WAHIS reports uses both semi-structured (.pdf file organized into sections, including free text) and structured data (.xlsx format) to display the reports. Each selected report underwent manual review by a veterinarian, guaranteeing a full understanding of the content and context. Information was manually extracted and hand-coded.

The following event information were extracted and entered into a structured template within a dedicated .csv file:

Named entities (e.g. animal species, country, city, variant);
Dates: when the case was laboratory confirmed, reported by OIE-WAHIS, and published;
Metrics: number of cases, number of deaths, number of susceptible animals.

Moreover, the following information on animal patient(s)/case(s) were extracted to populate the dataset:

Age;
Sex;
Living conditions;
Main reason for testing;
Suspected source of infection;
Symptoms: main reported clinical signs allegedly associated to SARS-CoV-2 were summarized with one to several keywords mentioned in the text. Multiple symptoms were separated by the operator “and”.

Extracted data described above was entered into the dataset as mentioned in the report and no information was subjected to any interpretation before entry. In addition, to facilitate understanding of the data and analysis, we have added the two following patient attributes:

The scientific name of the animal species was extracted from the report or inferred from the mentioned common name (when possible);
The family to which the animal species belongs was retrieved from expert knowledge or the literature.

Finally, for each SARS-CoV-2 event recorded in the dataset, we reported the primary and secondary source of information, i.e. source name (ProMED-mail or OIE-WAHIS) and link to the online report, as well as the original information source as referred by the primary source. A copy of each report used during the data extraction process was downloaded and saved as a PDF file. We inserted a timestamp on the saved file (ProMED-mail reports) or the download date was specified within the file name (it was not possible to insert a timestamp on OIE-WAHIS reports).

Data documenting each event corresponds to information available in the ProMED-mail and/or OIE-WAHIS report when consulting the report (see timestamp or download date). Potential subsequent editions or modifications of the report by ProMED-mail and/or OIE-WAHIS was not considered.

## Data Records

Each row of the dataset represents a SARS-CoV-2 event in animal(s), identified by a unique identifier (field **ID**). We consider as an event when one single case or several epidemiologically related cases were identified by the presence of viral RNA (proof of infection) and/or antibodies (proof of exposure) in an animal. Epidemiologically related cases include e.g. animals belonging to the same farm, captive animals housed together, pets belonging to the same household, or animals sampled within the same (generally transversal) study, featuring similar event and patient attributes, i.e. they underwent the same laboratory test(s) and showed the same results (including variant), exhibited the same symptoms and disease outcome, and were confirmed, reported (when applicable), and published on the same date (e.g. when pets of the same species sharing the same household showed different symptoms, they are reported as two distinct events). Events include follow-up history reports of outbreaks (e.g. follow-up on the clinical status of the animal, variant identification after case confirmation).

Each unique SARS-CoV-2 event is characterized by 46 quantitative and qualitative event and patient attributes (columns) that structure the dataset. Therefore, the dataset comprises five numeric fields (**number_cases**, **number_susceptible**, **number_tested**, **number_deaths**, and **age**), three date fields (**date_confirmed**, **date_reported**, and **date_published**), one character field (**sex**), and 29 string data fields, including the event unique identifier (**ID)** and two data fields requiring pre-defined string values (**epidemiological_unit** and **related_to_other_entries**). Eight metadata fields are dedicated to the information sources (e.g. names, links).

The field **related_to_other_entries** specifies potential relationship between events, and thereby allows identifying events that are related in space or time as well as follow-up reports (e.g. when animals described in two or more events are *living together* or when a follow-up report presents an *update of* another event, itself referred as *updated by*). Table 1 describes the 46 fields presented in the final dataset and their format. Supplementary File 1 provides three examples illustrating the structure and coding scheme of the SARS-ANI Dataset.

**Table 1.**
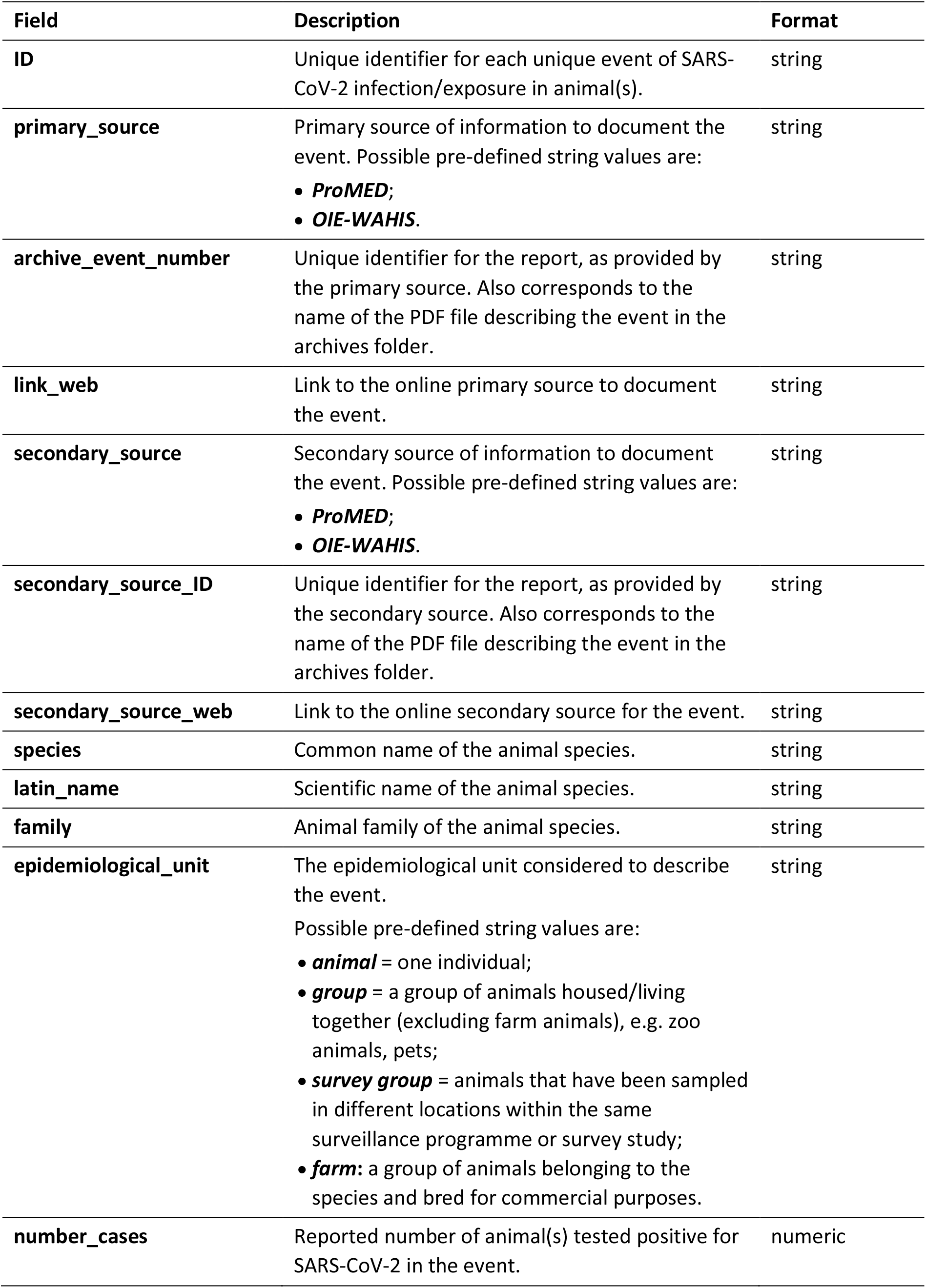
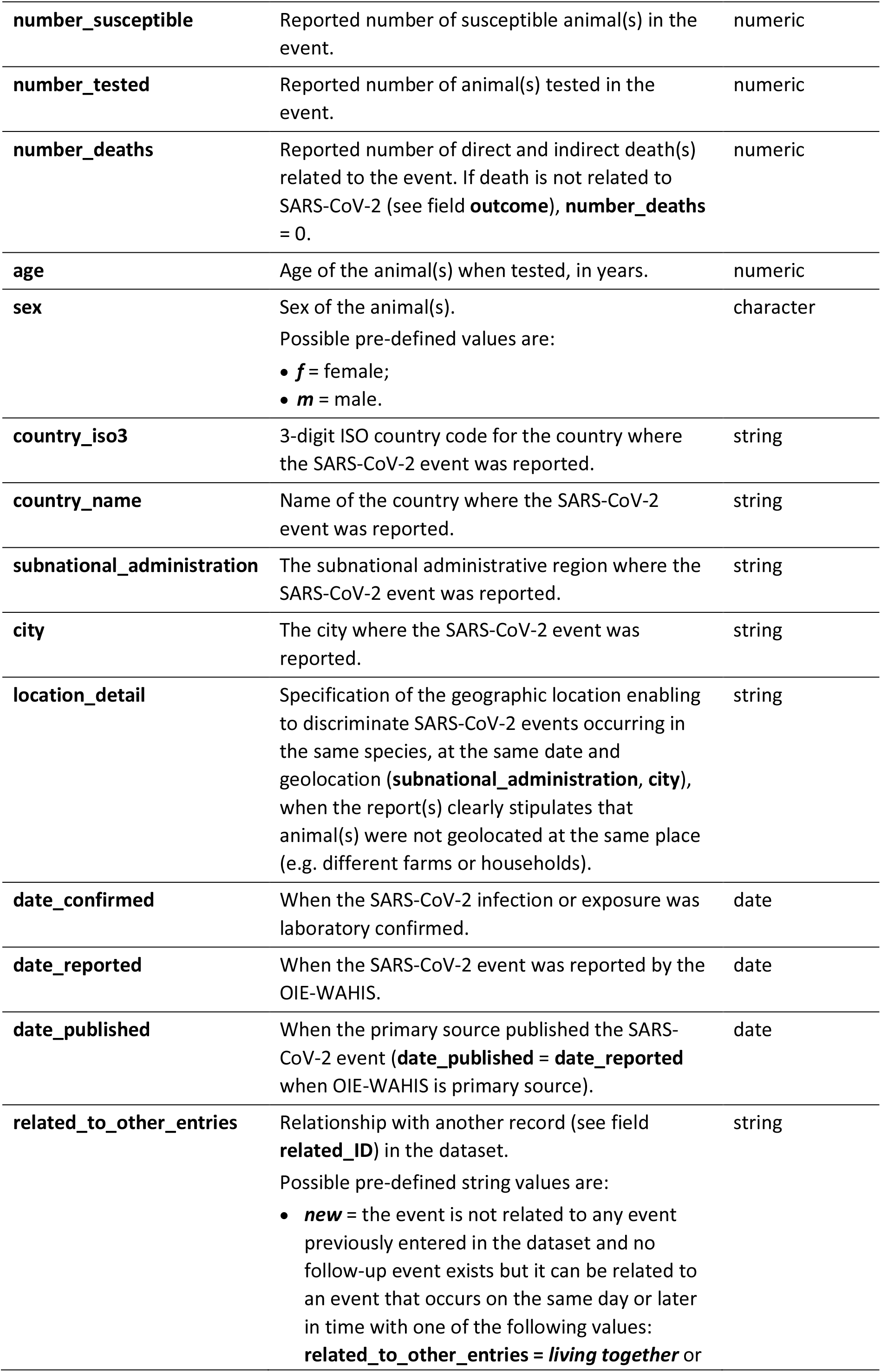
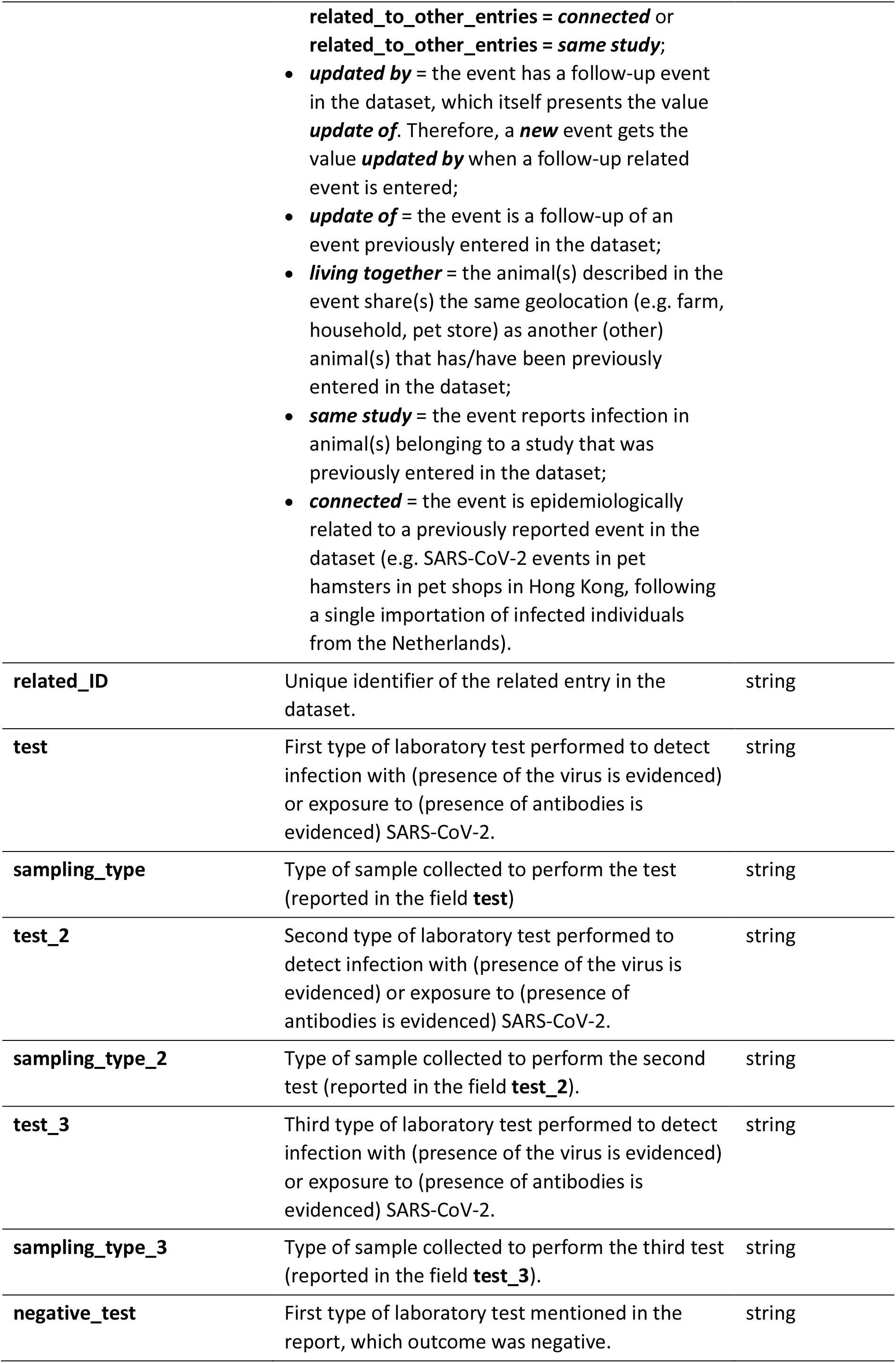
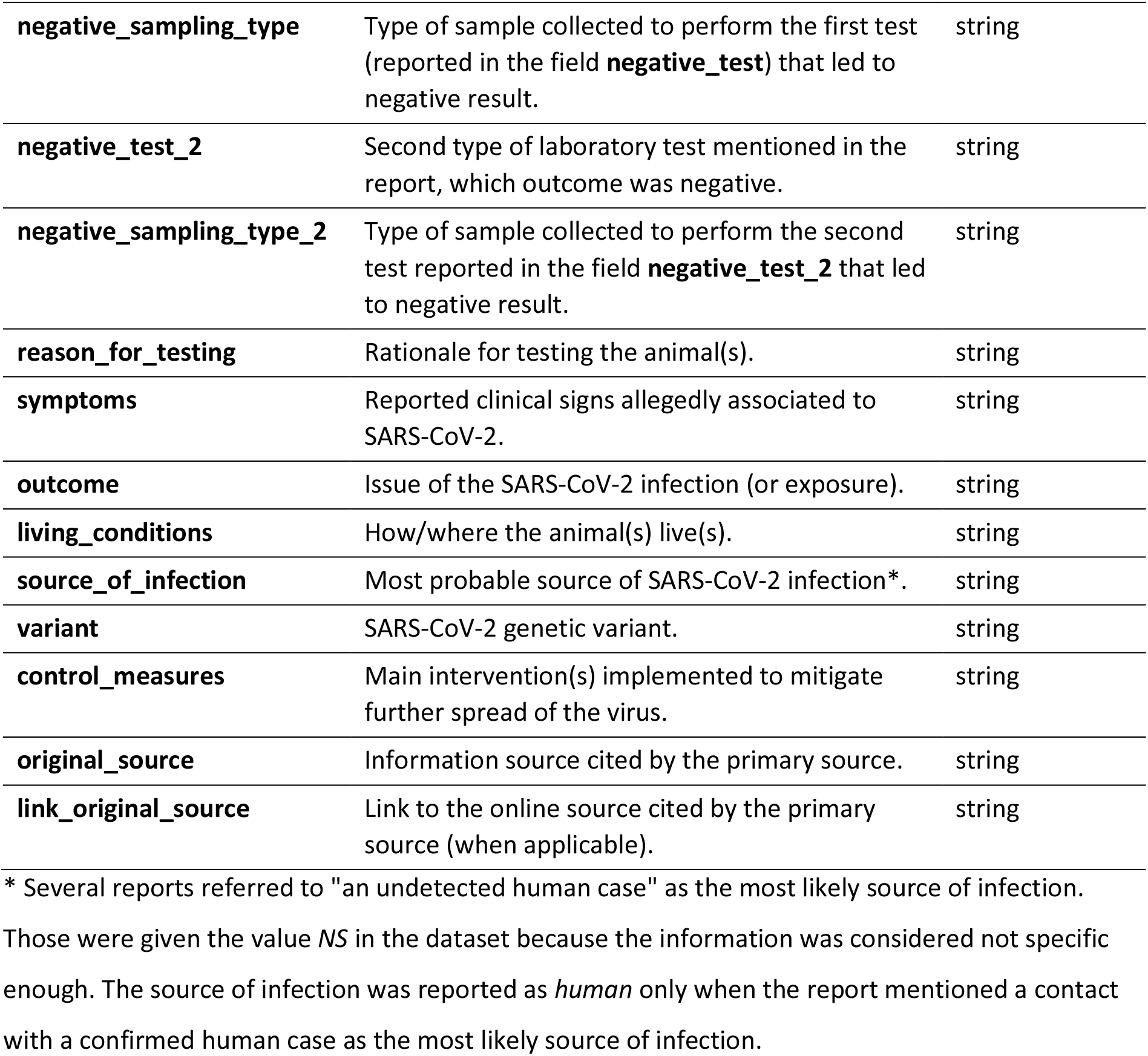
Description of the fields presented in the final dataset and format.

We have considered the two following values throughout the dataset:

*NA* (not applicable): when the field does not apply to the event. For example when only one laboratory test is conducted (field **test**), *NA* is reported for the second and third tests (**test_2** and **test_3**, respectively).
*NS* (not specified): when the information is relevant for the event but has not been specified in the report(s). For example, when a PCR test is performed for diagnostic purpose but the report(s) does not mention which sample was used, the sample type (**sampling_type**) is *NS*.

Accompanying the dataset, we publish the R code to perform technical and visual validation of the data as well as the ProMED-mail and OIE-WAHIS reports used as information sources (n = 353 reports as of date of submission). We also share the list of ProMED-mail and OIE-WAHIS reports that have not been included in the dataset and main reason for exclusion. This strategy, in line with the Open Science Principles^32^, aims to ensure that the data being reported is accurate and that all information can be accessed by researchers, policymakers, and the public. This also guarantees reproducibility of the data collection, may motivate further external validation processes as well as a large re-use of the data. The SARS-ANI files and products are summarized in Table 2. Event displays are also freely available on the SARS-ANI dashboard (https://vis.csh.ac.at/sars-ani/, see Usage Notes).

**Table 2.**
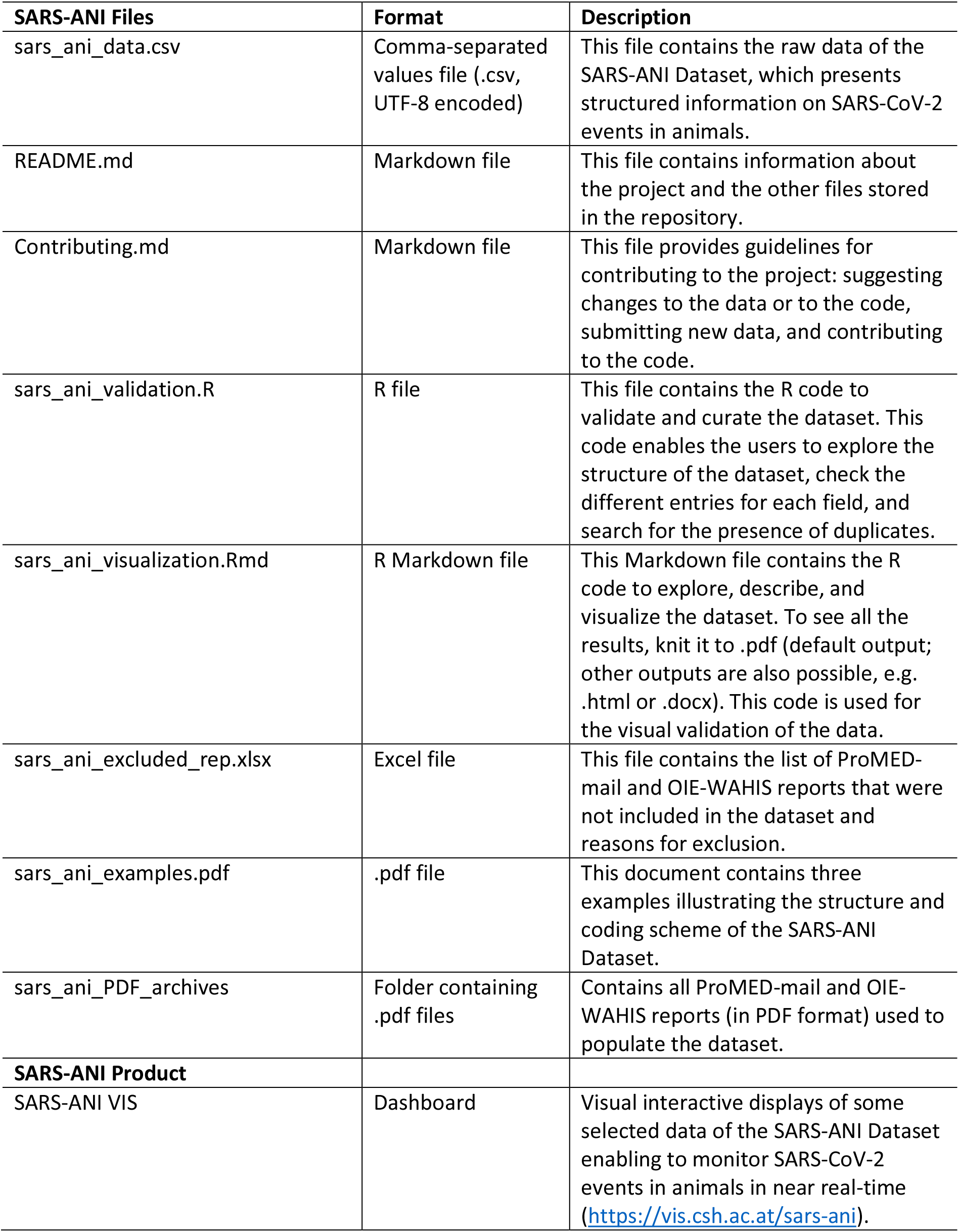
Details of the SARS-ANI files and products.

### Static Dataset

A static copy of the dataset in .csv file format is deposited on Zenodo^36^ (uploaded on 11 April 2022) and is publicly available at: https://doi.org/10.5281/zenodo.6442731, together with the SARS-ANI related files (metadata, R code, archived reports) described in Table 2. This static copy includes all SARS-CoV-2 events reported as of time of submission, spanning the period 26 February 2020 to 29 March 2022.

As of date of submission (11 April 2022), the SARS-ANI Dataset included 671 records of SARS-CoV-2 events in animals, representing 1,908 documented cases (infections and/or exposures), in 21 farmed, captive, wild, and domestic animal species belonging to 11 families, from 38 countries worldwide (covering 147 subnational administrative areas). The number of cases was not reported by ProMED-mail and/or OIE-WAHIS in 115 events. Fig. 2 shows the geographic distribution of the reported SARS-CoV-2 outbreaks included in the dataset. Table 3 summarizes the number of SARS-CoV-2 cases reported globally in each species.

**Fig. 2.**
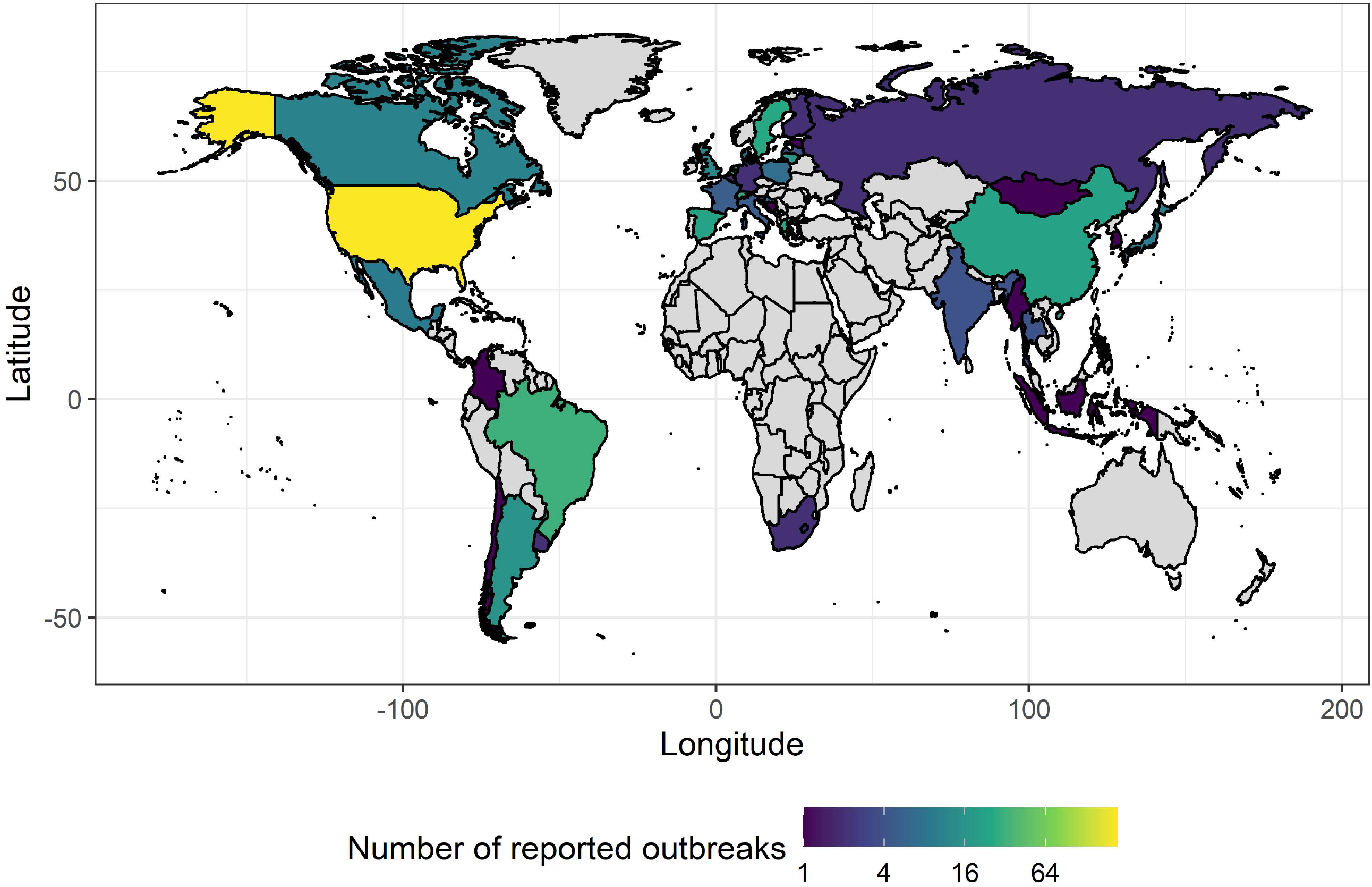
Geographic distribution of reported SARS-CoV-2 outbreaks (i.e. occurrence of one or more cases in an epidemiological unit) in animals per country. The number of outbreaks is lower than the number of events because distinct events i) may belong to the same epidemiological unit (e.g. animals that are living together, e.g. one farm, one household) or ii) may be follow-ups of the same outbreak. Note that if an outbreak is not published by ProMED-mail and/or OIE-WAHIS then it is not included in the dataset. Grey colour: no outbreak reported.

**Table 3.** Number of globally reported SARS-CoV-2 cases (infections or exposures) per animal species. (as of date of submission, 11 April 2022). This table includes only events for which the number of cases is documented.

| Family | Scientific name | Common name | Number cases |
| --- | --- | --- | --- |
| Mustelidae | *Neovison vison* | mink | 698* |
| Cervidae | *Odocoileus virginianus* | white-tailed deer | 433* |
| Felidae | *Felis catus* | cat | 323 |
| Canidae | *Canis familiaris* | dog | 201 |
| Felidae | *Panthera leo* | lion | 77 |
| Felidae | *Panthera tigris* | tiger | 75 |
| Hominidae | *Gorilla gorilla* | gorilla | 31 |
| Cricetidae | *Mesocricetus auratus* | hamster | 15 |
| Felidae | *Panthera uncia* | snow leopard | 14 |
| Mustelidae | *Mustela furo* | ferret | 9 |
| Mustelidae | *Aonyx cinereus* | otter | 8 |
| Castoridae | *Castor fiber* | beaver | 7 |
| Cricetidae | NS** | hamster | 3 |
| Felidae | *Puma concolor* | puma | 3 |
| Felidae | *Lynx canadensis* | lynx | 2 |
| Hippopotamidae | *Hippopotamus amphibius* | hippopotamus | 2 |
| Hyaenidae | *Crocuta crocuta* | hyena | 2 |
| Cervidae | *Odocoileus hemionus* | mule deer | 1 |
| Felidae | *Panthera pardus* | leopard | 1 |
| Felidae | *Prionailurus viverrinus* | fishing cat | 1 |
| Procyonidae | *Nasua nasua* | coatimundi | 1 |
| Viverridae | *Arctictis binturong* | binturong | 1 |

\* Number of cases was reported inconsistently in mink (data on the number of cases in mink is missing most of the time). Therefore, the number of diagnosed cases is largely under-estimated in mink. This is also true for deer, but to a lesser extent.
\*\* NS: Not specified in the reports. The hamster species was neither specified in the ProMED-mail nor OIE-WAHIS report.

### Dynamic Dataset

A live version of the SARS-ANI Dataset in .csv file format is publicly available on the GitHub repository, accessible at: https://github.com/amel-github/sars-ani, together with the related SARS-ANI files described in Table 2. We plan to update the dataset weekly for at least the next 12 months. The same technical procedures for data extraction and validation will be applied to any new event added to the dataset.

The GitHub interface allows users to flag potential inaccurate records in the dataset, which will trigger an error correction scheme, mainly consisting in processing the flagged record through another validation loop to check and replace the erroneous field when needed. Through the SARS-ANI GitHub repository and dashboard (https://vis.csh.ac.at/sars-ani/, see Usage Notes), we also expect to motivate experts in animal health, epidemiology, and conservation to support us filling up potential gaps in the records. The GitHub interface allows anybody to suggest changes to the dataset and code via the Issues Tracker (e.g. reporting an error and submitting new data). Contribution to the code can be implemented via a pull request (see Contributing.md, Table 2).

### Limitations

The dataset includes only SARS-CoV-2 events that have been published in ProMED-mail and/or OIE-WAHIS. Therefore, the integration of an event in the dataset strongly depends on the reporting strategy of the country to the OIE, the intensity of the research and surveillance strategy in the different animal species (e.g. whether pets from infected households are systematically investigated or not), the media coverage on the diagnosed cases, and the uptake of the reported event by the ProMED-mail team.

Moreover, we have identified five minor limitations in the dataset:

1. Some SARS-CoV-2 events in animals were not depicted in detail across the information sources, especially those related to infections in mink farms, for which the number of confirmed cases and deaths is often missing. Additionally, it was not always possible to discern from the report whether mink belonged to the same farm unit or not. In several reports on SARS-CoV-2 infections in mink farms, the number of sampled animals is specified whereas the number of positive animals is not (e.g. https://wahis.oie.int/#/report-info?reportId=16733), which make it impossible to infer whether the samples were pooled or not. Furthermore, SARS-CoV-2 infections in mink from Denmark and the Netherlands that occurred before 12 November 2020 (https://www.oie.int/en/oie-statement-on-covid-19-and-mink/) do not appear on the OIE-WAHIS interface (https://wahis.oie.int/#/events) while both countries endured a considerable burden in the fur-farming system^37^. Therefore, missing values (e.g. number of cases) for those events, reported in ProMED-mail, could not be completed. For these reasons, the dataset does not allow an accurate estimation of the economic and health burden of SARS-CoV-2 in mink.
2. Reports on mink and white-tailed deer often report the number of dams (sometimes dams and young) as the number of susceptible animals in the farm or herd, omitting adult males. We have reported the number as given in the information source, therefore, for these species, the number of susceptible animals in each event may be underestimated.
3. Although very seldom, some errors were found in the ProMED-mail or OIE-WAHIS reports. For example, when date of laboratory results precedes date of sampling or when sequencing is mentioned as first diagnostic test, although a PCR was most likely performed beforehand. Since we did not have any mean to retrieve the correct information, we entered the data as mentioned in the reports.
4. To respect and protect the privacy of the animal owners, outbreak location (i.e. city or village), as provided in the ProMED-mail and OIE-WAHIS reports, may not represent the exact location of the outbreak and should be interpreted with caution.
5. When multiple events are related (e.g. animals were living together: **related_to_other_entries** = *living together*), the number of susceptible animals (**number_susceptible**) from the same species is identical in both events and is therefore redundant. Similarly, when animals belonging to the same species and living together exhibited different symptoms or underwent different laboratory tests (therefore reported as distinct events) but were culled as part of a control strategy (i.e. **related_to_other_entries** = *living together* AND **control_measures** = *culling* OR *selective culling*), the number of reported deaths (**number_deaths**) is identical for these events (and is therefore redundant). Although very few records correspond to those cases, this may lead to a certain degree of over-counting the number of susceptible or dead animals if filtering of the events on the fields **related_to_other_entries** and **control_measures** is not performed accordingly (e.g. number of deaths should be counted once for farm animals living together). The code provided to explore the dataset presents examples of how using filters.

## Technical Validation

Validation of the collected data followed several steps (Fig. 1).

### Quality Control & Data Cleaning

First, the data underwent a quality control and cleaning procedures where the unique value of each field was checked to search for inaccurate (e.g. containing typographical errors or not belonging to a pre-defined list of entities), unreliable (e.g. the value was not specific), incorrectly formatted (e.g. date was formatted as dd/mm/yyyy instead of yyyy-mm-dd), or missing data in the dataset. This step was performed in R^38^ using the base function *unique(*). Events containing detected errors were manually inspected against original reports. When necessary, the erroneous values were modified, replaced, or removed.

### Search for Duplicate Events

We have identified duplicate events, defined as unique event reported more than once in the dataset. Events were flagged as duplicate if the geolocation information (i.e. **country**, **subnational_administration**, **city**, and **location_detail**), species denomination (**species** and **latin_name**), sex, age, symptoms, date (**date_confirmed** or **date_reported** if **date_confirmed** was missing or **date_published** if the two other dates were missing), number of cases, number of deaths, number of susceptible, tests conducted, outcome, and relationship to another event (**related_ID**) were identical. This step was executed in R^38^, using the dplyr package^39^. Duplicate events were flagged and manually inspected against original information source(s) to confirm redundancy. Duplicate events were removed. Events that were incorrectly flagged as duplicates were corrected. The code to reproduce the first and second steps of the technical validation into R^38^ (Table 2) is freely accessible at: https://github.com/amel-github/sars-ani and at https://doi.org/10.5281/zenodo.6442731.

### Visual Validation

The data was visually inspected through different graphical displays. Figures and maps were produced in R^38^, using the packages ggplot2^40^, webr^41^, and visNetwork^42^ for graphical visualizations. The code to visually summarize the data (through maps, figures, and interactive network) is provided as an R Markdown document (Table 2) and is publicly accessible at https://github.com/amel-github/sars-ani and at https://doi.org/10.5281/zenodo.6442731.

### Expert Validation

“*Expert judgment is defined as an informed opinion of people with experience in the subject, who are recognized by others as qualified experts in it, and who can provide information, evidence, judgments and assessments*”^43^. First, the project leader reviewed unresolved issues met during data collection, conducted random verification of the events recorded in the dataset against the original reports, and randomly checked entry of ProMED-mail and OIE-WAHIS reports in the dataset. In a second step, two experts in our team examined the data as well as its graphical displays and subsequently provided feedback. These expert validation steps enabled to clarify questions and further resolve potential omissions in the dataset.

## Usage Notes

The SARS-ANI Dataset is currently the most comprehensive global dataset of SARS-CoV-2 events reported in animals. Information displays in the dataset aim to facilitate a wide range of analyses of SARS-CoV-2 infections/exposures in animals that will further our understanding of the epidemiology and impact of SARS-CoV-2 in the different animal species, at different scales: international, national, or subnational. The standardized coding of the health events makes the dataset intelligible (also for non-experts) while the data format allows for a great analytical flexibility and interesting integration potential. We hope that these qualities will enhance the reuse and combination of the data across sectors and disciplines.

### Addressing SARS-CoV-2 Events in Animals

Fig. 3, Fig. 4, and Fig. 5 visually display answers to some of the many questions that can be investigated using the data; other questions/visualizations can be computed from the published code (https://github.com/amel-github/sars-ani and https://doi.org/10.5281/zenodo.6442731):

What is the case fatality rate of SARS-CoV-2 per animal species and country? (Fig. 3)
Which SARS-CoV-2 variants have been identified in the different animal species? (Fig. 4)
Why were animals tested for SARS-CoV-2? (Fig. 5)

**Fig. 3.**
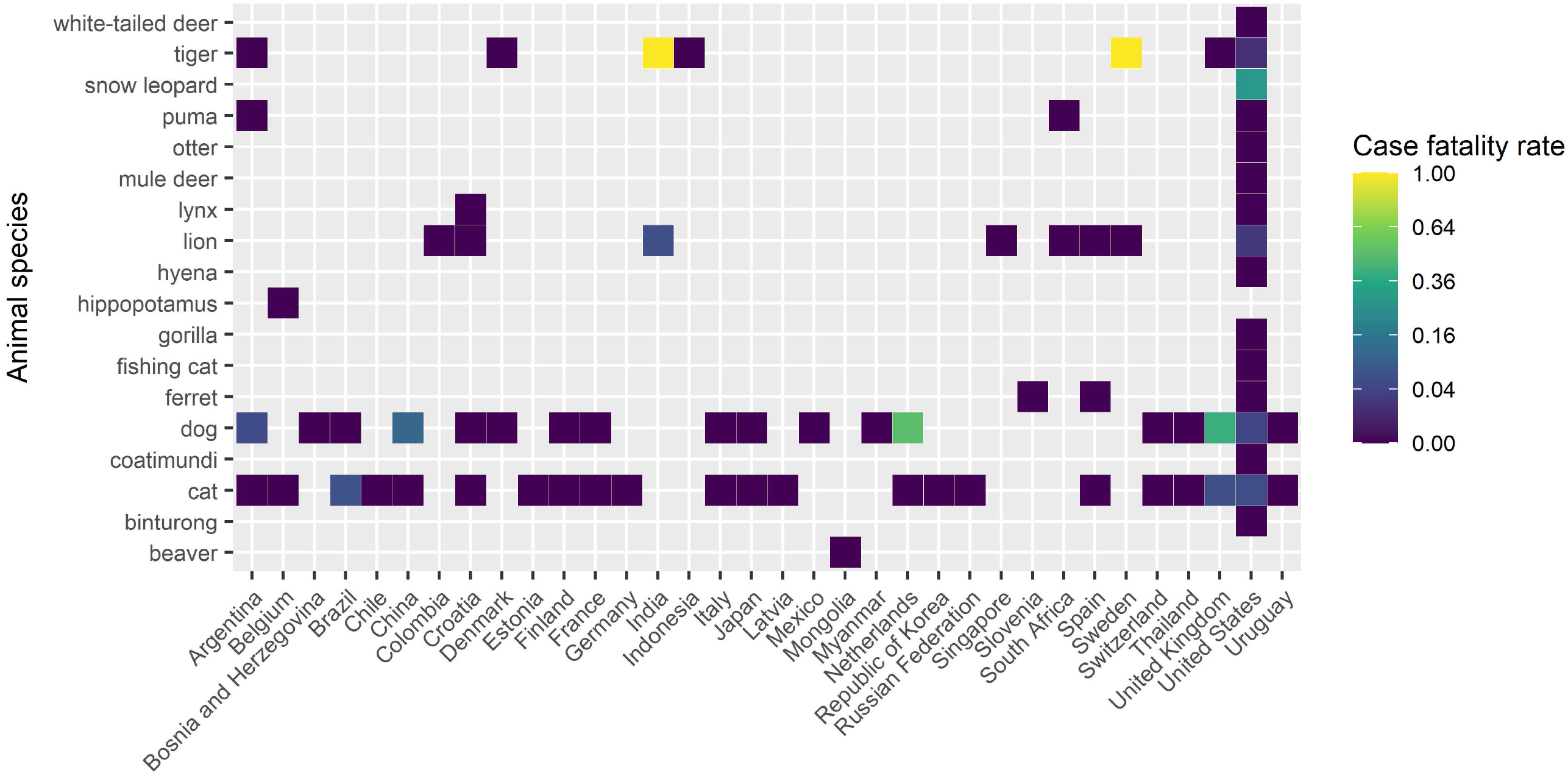
SARS-CoV-2 case fatality rate (CFR) per species and country. The CFR for each species and country is obtained by dividing the total number of reported deaths in one species by the total number of reported cases for this species in the country. Animals culled as part of a control strategy are excluded (not all were diagnosed as infected). Similarly, mink are not included here because data on case and death numbers are partial. The CFR depends strongly on testing and does not give information on the infection fatality rate (IFR, number of deaths divided by the total number of infected individuals) or mortality rate (MR, number of deaths divided by the total at-risk population).

**Fig. 4.**
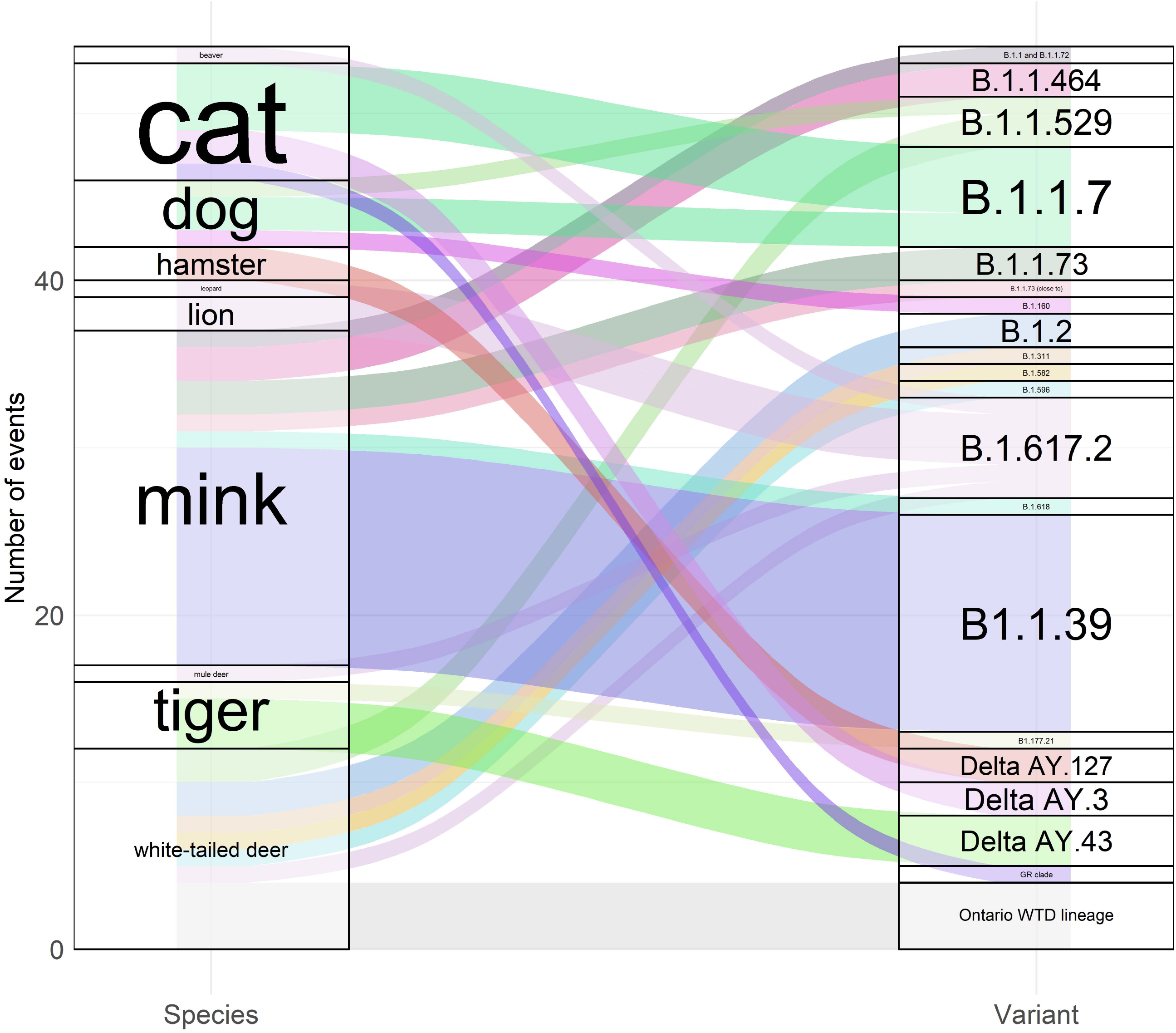
Sankey diagram showing the SARS-CoV-2 variants identified in the different animal species. The figure describes the number of events (one event may include one or more cases).

**Fig. 5.**
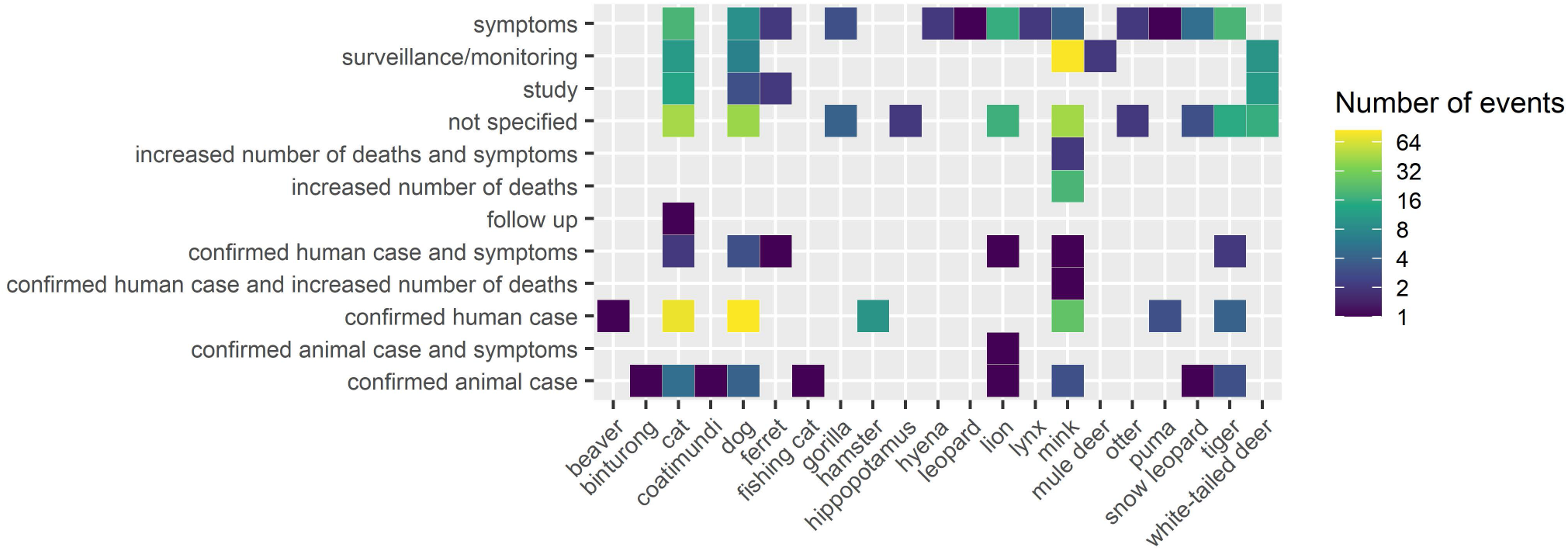
Rationales for testing animals for SARS-CoV-2 infection or exposure. Only positive animals are reported in the dataset; investigations that led to negative results are not (or rarely) reported to the authorities or media.

The dataset can become an essential tool to estimate the threat and impact of SARS-CoV-2 on animal husbandry (specifically mink/fur animals), pets, wildlife (including captive wild animals), and conservation programmes^29,44^. Coupled with economic data on the cost of testing, implemented control measures (e.g. mass culling) and value of farmed and captive animals, it can facilitate research in animal health economics, e.g. assessing the economic burden of SARS-CoV-2 infections on animal production systems and conservation programmes or supporting cost-benefit analysis of prevention of zoonotic-origin pandemics^45^. The dataset can also assist risk-based veterinary surveillance by identifying surveillance needs to protect animal health and efficiently prioritizing resource allocation, especially in resource-limited contexts.

Additionally, the SARS-ANI data can be ported and integrated into existing platforms on animal diseases (e.g. the Wildlife Health Information Sharing Partnership: https://whispers.usgs.gov/home or the WildHealthNet initiative: https://oneworldonehealth.wcs.org/Initiatives/WildHealthNet.aspx). Finally, the procedure and the standardized reporting format developed in SARS-ANI are applicable to other infectious threats; likewise, the flexible and well-documented analytical tools can be adapted and used for descriptive epidemiology of other diseases.

### One Health Surveillance Strategy

Because surveillance of zoonotic-origin diseases at the human-animal-environment interface is extremely challenging, there is a need for comprehensive One Health approaches to monitor SARS-CoV-2 carriage and infections in animals globally^18^. One Health tools that enable the integrative analysis and visualization of SARS-CoV-2 events are critical. The SARS-ANI Dataset can be combined with other data across sectors (e.g. data on COVID-19 cases in humans, land-use and environmental data) to support research efforts on SARS-CoV-2 at the human-animal-environment interface^46^, e.g. identifying hotspots of circulation and spillover, developing and adapting integrated One Health surveillance systems of SARS-CoV-2 events, and elucidating the natural ecology of the SARS-CoV-2. We believe the SARS-ANI Dataset, with timely and reliable information, can assist inter-professional and multi-sectoral SARS-CoV-2 prevention and control activities, including the development of relevant national and international regulations and agreements to improve preparedness and reduce the risk of transmission between humans and animals^47^. Information sharing on SARS-CoV-2 infections/exposures in animal living close to humans will also benefit veterinary and public health professionals in their investigations of SARS-CoV-2 cases at the human-animal interfaces (e.g. need for testing pets/captive animals in contact with COVID-19 diagnosed owners/caretakers; types of samples and tests to be performed). Furthermore, the data can provide practical advice for the international trade in domestic and farmed species, for which a role in human infections has been recognized^10^. Finally, the data can be used to develop and adapt national or global One Health prevention, preparedness and response plans for emerging coronavirus diseases and assist public health officers in their task.

### SARS-ANI VIS: Informing Scientists, Stakeholders, and the Public

The SARS-ANI dashboard, publicly accessible at: https://vis.csh.ac.at/sars-ani/, provides intuitive insights into specific aspects of SARS-CoV-2 events in animals at-a-glance (Fig. 6). The visual representations of the data are displayed in a narrative format, where information is conveyed through vertically connected segments. Each segment features a selected topic, starting from a general overview and leading to more specific questions, such as variants across species or reported clinical signs. The partition into segments aims to ease the understanding of the information and facilitate webpage navigation. Each segment is carefully designed to exhibit the intended information in a fashion that can be comprehended by scientists and the general public alike. The dashboard thus will facilitate access to the data, favour animal health information sharing, and foster global understanding of the data among the scientific community, stakeholders, and the public. The dashboard intends to support public education about the risk of SARS-CoV-2 transmission between humans and animals and raise public awareness about possible wildlife conservation issues posed by the SARS-CoV-2 pandemic. The dashboard is linked to the live dataset available on GitHub (https://github.com/amel-github/sars-ani) and will therefore be subjected to continuous updates.

**Fig. 6.**
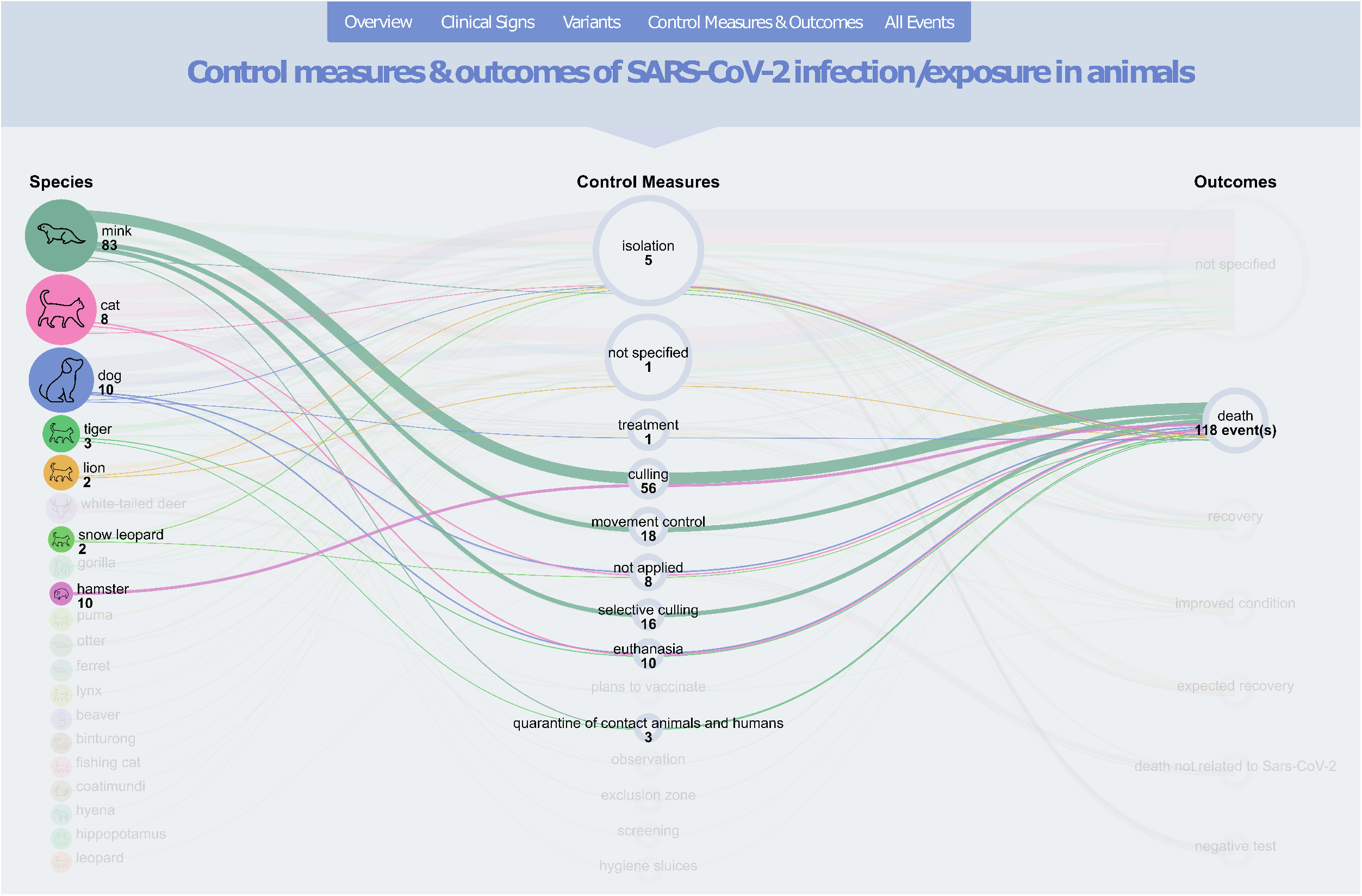
Screen shot of the SARS-ANI dashboard.

## Code Availability

A static version of the SARS-ANI Dataset and related files that accompany the dataset (metadata, R code, archived reports) have been deposited on Zenodo^36^ and are available at: https://doi.org/10.5281/zenodo.6442731, for public access. A live version of the dataset (together with the related files) is accessible on GitHub at: https://github.com/amel-github/sars-ani. Please refer to the README file in the code release for further instructions.

## Acknowledgements

The authors thank their colleagues of the Unit Veterinary Public Health and Epidemiology, University of Veterinary Medicine Vienna, for their inputs on the different visual displays of the data. The authors thank W. Schueller for his support with the GitHub interface.

## Author contributions

A.N. developed the structured template and the coding scheme, extracted the data, investigated flagged errors and redundancies, updated the dataset, archived the information sources, and helped write the manuscript.

A.D.L. conceived the study, coordinated the production of the dataset, developed the code for technical and visual validation, created the figures and tables, and helped write the manuscript.

L.Y. created the dashboard, including conceptualisation, developing the data story and data automation process.

J.S. supported the development of the dashboard and helped write the manuscript.

A.K. and C.W. contributed to the data review process and helped write the manuscript.

## Supplementary File 1

Below are three examples illustrating the structure and coding scheme of the SARS-ANI Dataset. These examples intend to facilitate the comprehension of the data as well as the coding of potential relationships between events.

### Example 1

The two following SARS-CoV-2 animal events occurred in the United States (**country_name**), in one household (**living_conditions =** *pet* AND event *2* is related to event 1, see **related_to_other_entries** = “living together”) where three dogs were living (**number_susceptible =** *3*). The first dog was tested by PCR (**test =** *PCR*) because it had contact with a human diagnosed with COVID-19 (**reason_for_testing =** *confirmed human case*); the infection was confirmed on 2020-06-25 (**date_confirmed**): the dog showed symptoms that were not related to SARS-CoV-2 infection (**symptoms =** *unrelated symptoms*) and died from a cause not related to its SARS-CoV-2 status (**outcome** = *death not related to Sars-CoV-2*). This case was reported (**date_reported**) on 2020-07-02 by the OIE-WAHIS.

Almost two months later (**date_reported** = *2020-08-27*), a second dog of this household (counting at that time two dogs, **number_susceptible** = *2*) which had contact with the dog described in event 1 (**related_to_other_entries** = *living together* AND **reason_for_testing =** *confirmed animal case*) was tested for SARS-CoV-2 by virus neutralization test (**test**). The dog was asymptomatic (**symptoms =** *subclinical*) and was isolated (**control_measures**). Outcome of the infection was not reported (outcome = *NS*).

#### Summary

These events describe one outbreak of SARS-CoV-2 in a multi-dog household following contact with a COVID-19-infected person and in which 2/3 dogs were infected (for clarity purpose, not all fields of the dataset are shown below).

| ID | species | epidemiological_unit | number_cases | number_susceptible | number_tested | number_deaths |
| --- | --- | --- | --- | --- | --- | --- |
| event 1 | dog | animal | 1 | 3 | 2 | 0 |
| event 2 | dog | animal | 1 | 2 | 1 | 0 |

| country_name | date_confirmed | date_reported | date_published | related_to_other_entries | related_ID | test |
| --- | --- | --- | --- | --- | --- | --- |
| United States | 2020-06-25 | 2020-07-02 | 2020-07-03 | new | NA | PCR |
| United States | NS | 2020-08-27 | 2020-08-27 | living together | event 1 | virus neutralisation test |

| sampling_type | reason_for_testing | symptoms | outcome | living_conditions | source_of_infection | control_measures |
| --- | --- | --- | --- | --- | --- | --- |
| NS | confirmed human case | unrelated symptoms | death not related to Sars-CoV-2 | pet | human | NA |
| NS | confirmed animal case | subclinical | NS | pet | animal | isolation |

### Example 2

The three following events occurred in a zoo (**living_conditions)** in the United States (**country_name**). A tiger living with four other animals from the same species (**number_susceptible** = 1+4 = *5*) was confirmed positive with SARS-CoV-2 by PCR (**test** = *PCR*) on 2020-04-03. Event 1 described infection in one animal (**epidemiological_unit** = *animal*). Date of reporting (**date_reported**) and publishing (**date_published**) are similar (*2020-04-06*). Condition of this tiger improved (**outcome** = *improved condition*).

Later in the month, two other events were reported in which SARS-CoV-2 infection was diagnosed by PCR (**test** = *PCR*) in the four other animals living with the tiger reported in event 3 (**related_to_other_entries** = *living together*). For both events, **date_confirmed** is not specified (*NS*), **date_reported** = *2020-04-17* and **date_published** = *2022-05-25*. For one animal (event 4), no information is available regarding the outcome of the infection (**outcome** = *NS*). For the three other animals described in event 5, the outcome of the infection is reported (**outcome** = *improved condition*). Because these three tigers were reported together and showed the same event and patients attributes (e.g. **date_reported**, **test**, **outcome**), they are entered as one event in the dataset (**epidemiological_unit** = *group*).

#### Summary

These three events describe one outbreak (occurrence of one or more cases in an epidemiological unit, which is here the enclosure of the tigers in the zoo), which started on 2020-04-03 and included five cases of SARS-CoV-2 infection (for clarity purpose, not all fields of the dataset are shown below).

| ID | species | epidemiological_unit | number_cases | number_susceptible | number_tested |
| --- | --- | --- | --- | --- | --- |
| event 3 | tiger | animal | 1 | 5 | 1 |
| event 4 | tiger | animal | 1 | 5 | NS |
| event 5 | tiger | group | 3 | 5 | NS |

| number_deaths | country_name | date_confirmed | date_reported | date_published | related_to_other_entries |
| --- | --- | --- | --- | --- | --- |
| 0 | United States | 2020-04-03 | 2020-04-06 | 2020-04-06 | new |
| 0 | United States | NS | 2020-04-17 | 2020-04-25 | living together |
| 0 | United States | NS | 2020-04-17 | 2020-04-25 | living together |

| related_ID | test | outcome | living_conditions | source_of_infection | variant |
| --- | --- | --- | --- | --- | --- |
| NA | PCR | improved condition | zoo | NS | NA |
| event 3 | PCR | NS | zoo | NS | NA |
| event 3 | PCR | improved condition | zoo | NS | NA |

### Example 3

The three following events report an outbreak that occurred in free ranging (**living_conditions** = *wildlife*) white-tailed deer (**species**) in Canada (**country**) that were tested during a surveillance/monitoring programme (**reason_for_testing**). The animals were sampled as part of the same epidemiological investigation but not necessary at the same geographic location (**epidemiological_unit** = *survey group*). Event 7 is an *update of* (**related_to_other_entries**) event 6 (**related_to_other_entries** = *update by*) and event 8 (related to event 7, see below) should also be considered as an *update of* event 6.

Events 6 and 7 report infections in more than one animal (**epidemiological_unit** = *survey group*) while event 8 reports infection in one single animal (**epidemiological_unit** = *animal*) related to the group described in event 7 (**related_to_other_entries** = *same study*). Event 7 and 8 are distinguished because the tests conducted in the animal in event 8 differed from those performed on the four animals described in event 7.

Date at which infection was confirmed (**date_confirmed)** is not specified in any events (*NS*); only event 6 was reported by the OIE-WAHIS (**date_reported** = *2022-01-20*). The follow-up events (events 7 and 8) were published ~1.5 month after event 6. Most likely, this difference is due to the time required for laboratory analyses (events 7 and 8 present more sample and test types and provide identification of the SARS-CoV-2 variant involved).

#### Summary

These three events describe one outbreak (occurrence of one or more cases in an epidemiological unit, which is here the surveyed animals) which was reported for the first time by OIE-WAHIS and published on 2022-01-20 and was subsequently updated on 2022-03-03 in ProMED-mail, probably following more laboratory investigations. The outbreak included five confirmed cases of SARS-CoV-2 in white-tailed deer in Canada. The five animals did not die following SARS-CoV-2 infection (**number_deaths** = *0* AND **outcome =** *death not related to Sars-CoV-2*). Indeed, they died from hunting, which this is not specified in the dataset but can be found in the reports (for clarity purpose, not all fields of the dataset are shown below).

| ID | species | epidemiological_unit | number_cases | number_tested | number_deaths |
| --- | --- | --- | --- | --- | --- |
| event 6 | white-tailed deer | survey group | 5 | 213 | 0 |
| event 7 | white-tailed deer | survey group | 4 | 213 | 0 |
| event 8 | white-tailed deer | animal | 1 | 213 | 0 |

| country_name | date_confirmed | date_reported | date_published | related_to_other_entries | related_ID |
| --- | --- | --- | --- | --- | --- |
| Canada | NS | 2022-01-20 | 2022-01-20 | updated by | event 7 |
| Canada | NS | NA | 2022-03-03 | update of | event 6 |
| Canada | NS | NA | 2022-03-03 | same study | event 7 |

| test | sampling_type | test_2 | sampling_type_2 | test_3 | sampling_type_3 |
| --- | --- | --- | --- | --- | --- |
| PCR | NS | NA | NA | NA | NA |
| PCR | lymph node (post mortem) | PCR | nasal swab | gene sequencing | nasal swab |
| PCR | nasal swab | gene sequencing | nasal swab | NA | NA |

| reason_for_testing | symptoms | outcome | living_conditions | variant |
| --- | --- | --- | --- | --- |
| surveillance/monitoring | subclinical | death not related to Sars-CoV-2 | wildlife | NA |
| surveillance/monitoring | subclinical | death not related to Sars-CoV-2 | wildlife | Ontario WTD lineage |
| surveillance/monitoring | subclinical | death not related to Sars-CoV-2 | wildlife | Ontario WTD lineage |

